# Inter-individual variability of neurotransmitter receptor and transporter density in the human brain

**DOI:** 10.1101/2025.09.09.674944

**Authors:** Justine Y. Hansen, Jouni Tuisku, Jarkko Johansson, Zeyu Chang, Colm J. McGinnity, Vincent Beliveau, Synthia Guimond, Melanie Ganz, Martin Nørgaard, Marian Galovic, Gleb Bezgin, Sylvia M. L. Cox, Jarmo Hietala, Marco Leyton, Eliane Kobayashi, Pedro Rosa-Neto, Thomas Funck, Nicola Palomero-Gallagher, Gitte M. Knudsen, Paul Marsden, Alexander Hammers, Lauri Nummenmaa, Lauri Tuominen, Bratislav Misic

**Affiliations:** Montréal Neurological Institute, McGill University, Montréal, QC, Canada; University of Turku and Turku University Hospital, Finland; Umeå University, Umeå, Sweden; King’s College London & Guy’s and St Thomas’ PET Centre, King’s College London, London, UK; Medical University of Innsbruck, Innsbruck, Austria; Neurobiology Research Unit, Copenhagen University Hospital Rigshospitalet, Copenhagen, Denmark; Department of Psychiatry, The Royal’s Institute of Mental Health Research, University of Ottawa, ON, Canada; Department of Psychoeducation and Psychology, University of Quebec in Outaouais, Gatineau, QC, Canada; Department of Computer Science, University of Copenhagen, Copenhagen, Denmark; Molecular Imaging Branch, National Institute of Mental Health (NIMH), USA; Clinical Neuroscience Center, University Hospital Zurich, Zurich, Switzerland; UCL Queen Square Institute of Neurology, London, UK; MRI Unit, Chalfont Centre for Epilepsy, UK; Department of Psychiatry, McGill University, Montréal, QC, Canada; Department of Neurology and Peter O’Donnell Jr. Brain Institute, University of Texas Southwestern Medical Center; Center for the Developing Brain, Child Mind Institute, New York, USA; Institute of Neuroscience and Medicine (INM-1), Research Centre Jülich, Jülich, Germany; C. and O. Vogt Institute for Brain Research, Medical Faculty, University Hospital Düsseldorf, Heinrich-Heine University Düsseldorf, Düsseldorf, Germany; Institute of Clinical Medicine, University of Copenhagen, Copenhagen, Denmark; Research Department of Biomedical Computing & Research Department of Early Life Imaging, King’s College London, School of Biomedical Engineering and Imaging Sciences

## Abstract

Neurotransmitter receptors guide the propagation of signals between brain regions. Mapping receptor distributions in the brain is therefore necessary for understanding how neurotransmitter systems mediate the link between brain structure and function. Normative receptor density can be estimated using group averages from Positron Emission Tomography (PET) imaging. However, the generalizability and reliability of group-average receptor maps depends on the inter-individual variability of receptor density, which is currently unknown. Here we collect group standard deviation brain maps of PET-estimated protein abundance for 12 different neurotransmitter receptors and transporters across 7 neurotransmitter systems, including dopamine, serotonin, acetylcholine, glutamate, GABA, cannabinoid, and opioid. We illustrate how cortical and subcortical inter-individual variability of receptor and transporter density varies across brain regions and across neurotransmitter systems. We complement inter-individual variability with inter-regional variability, and show that receptors that vary more across brain regions than across individuals also demonstrate greater out-of-sample spatial consistency. Altogether, this work quantifies how receptor systems vary in healthy individuals, and provides a means of assessing the generalizability of PET-derived receptor density quantification.

## INTRODUCTION

Neurotransmitter receptors modulate neuronal activity, guide synaptic wiring, and mediate brain-wide communication. Mapping neurotransmitter receptor distributions in the brain is therefore necessary for understanding how chemoarchitecture shapes brain structure and function. We recently collated a Positron Emission Tomography (PET) atlas of in vivo whole-brain neurotransmitter receptor and transporter densities across 19 unique receptors and transporters and 9 neurotransmitter systems [23, 37]. This atlas is widely used for studying chemoarchitectonic mechanisms underlying, for example, neural rhythms [60], pharmacological perturbations [34, 67], energy metabolism [11], cognition [72], and multiple diseases and disorders [24, 27, 41, 52, 71].

Nevertheless, brain anatomy and function vary across individuals, manifesting as individual differences in cognition and behaviour [9, 43, 59]. In addition, brain regions and systems develop at different rates, and are differentially subjected to influence from the environment (e.g. via sensory stimuli) and transcriptomic programs [10, 64]. Inter-individual variability in receptor density may therefore be greater in some brain regions than in others. Some inferences on the inter-individual variability of receptor density can be made from group-average receptor density maps alone: group receptor density brain maps can be compared across sites, PET tracers, imaging modalities, and even across biological features (e.g. receptor density versus protein-coding gene expression) [8, 21, 23, 44, 45]. However, these strategies can only assess the spatial similarity of brain maps rather than the inter-individual variability of regional receptor density.

To better understand how receptor abundance varies across individuals, we collate group standard deviation maps for 12 neurotransmitter receptors and transporters across 7 neurotransmitter systems and nearly 700 individuals. We show cortical and subcortical brain maps of inter-individual receptor abundance variability, and benchmark receptor variability across PET tracers. We then compare inter-individual and inter-regional variability. By interpreting the present findings alongside previous work comparing spatial distributions of receptors, we provide receptor-specific hypotheses for sources of variability. Altogether, this work serves as a reference point for assessing receptor and transporter measurement generalizability in the human brain.

## RESULTS

We collated group standard deviation maps of PET-derived neurotransmitter receptor and transporter densities from a total of 12 different receptors/transporters across 7 neurotransmitter systems, including dopamine, serotonin, acetylcholine, glutamate, GABA, cannabinoid, and opioid (Table 1). All mean and standard deviation maps are parcellated according to 100 cortical regions [57] and 54 subcortical regions [66] (note that allocortex (e.g. hippocampus) is included in the subcortical atlas). Given that standard deviations scale with the mean (Fig. S1, S2), we normalize standard deviation by the mean, resulting in a brain map of the within-region inter-individual coefficient of variation for each neurotransmitter receptor and transporter (Fig. 1, Fig. 2). In both cortex and subcortex, inter-individual coefficient of variation is heterogeneously distributed and highly organized across brain regions. For many receptors and transporters, cortical coefficient of variation appear greatest in unimodal brain regions, including primary somatomotor and somatosensory cortex as well as primary visual cortex (Fig. 1). Meanwhile, subcortical coefficient of variation is often greatest in ventral structures as well as the caudate (Fig. 2).

**TABLE 1.**
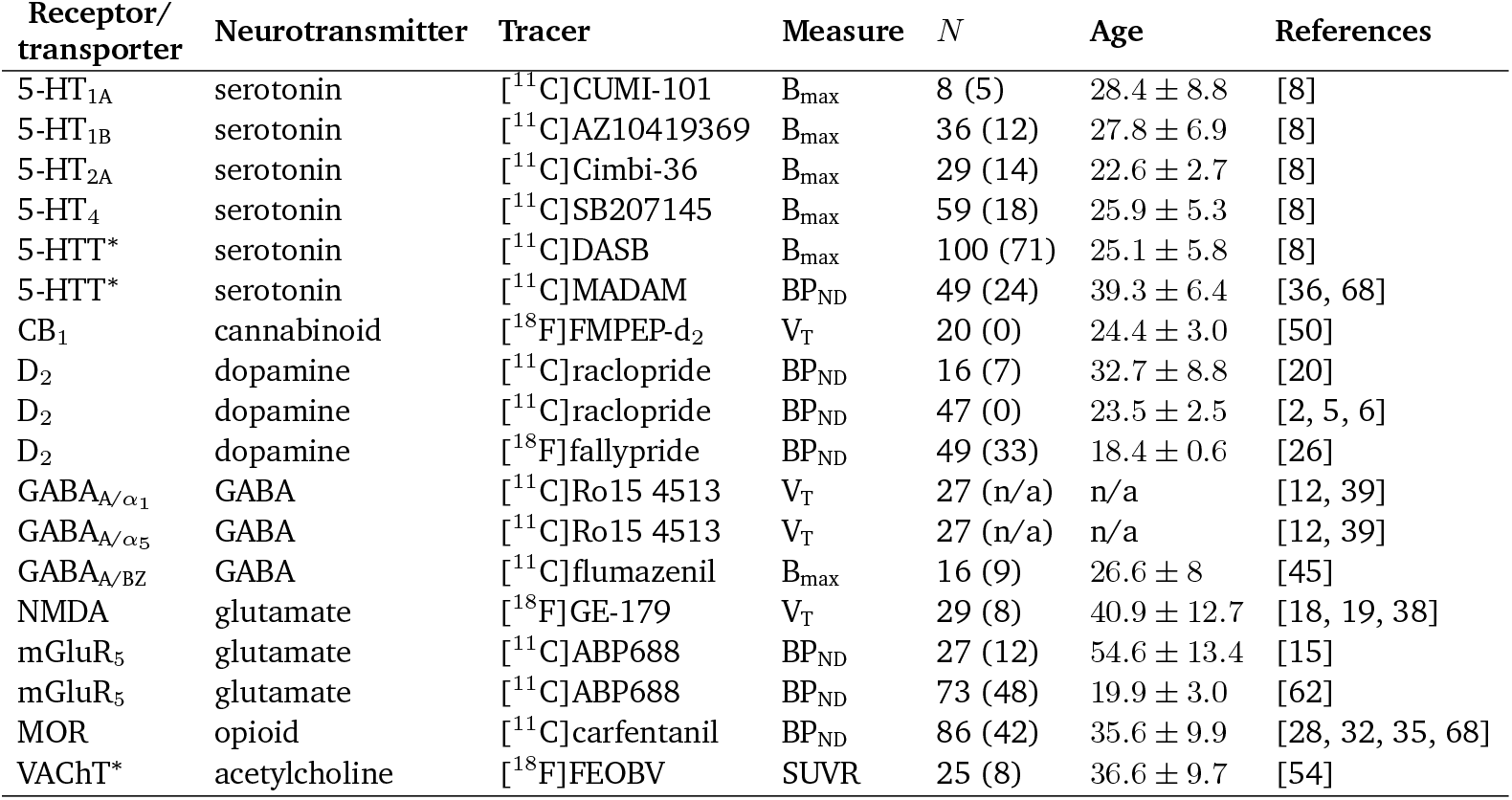
Neurotransmitter receptors and transporters included in analyses. BP_ND_ = non-displaceable binding potential; V_T_ = tracer distribution volume; B_max_ = density (pmol/ml) converted from binding potential using autoradiography-derived densities; SUVR = standard uptake value ratio. Values in parentheses (under *N*) indicate number of females. Asterisks indicate transporters.

| Receptor/<br>transporter | Neurotransmitter | Tracer | Measure | $N$ | Age | References |
| --- | --- | --- | --- | --- | --- | --- |
| 5-HT<sub>1A</sub> | serotonin | [<sup>11</sup>C]CUMI-101 | B<sub>max</sub> | 8 (5) | 28.4 ± 8.8 | [8] |
| 5-HT<sub>1B</sub> | serotonin | [<sup>11</sup>C]AZ10419369 | B<sub>max</sub> | 36 (12) | 27.8 ± 6.9 | [8] |
| 5-HT<sub>2A</sub> | serotonin | [<sup>11</sup>C]Cimbi-36 | B<sub>max</sub> | 29 (14) | 22.6 ± 2.7 | [8] |
| 5-HT<sub>4</sub> | serotonin | [<sup>11</sup>C]SB207145 | B<sub>max</sub> | 59 (18) | 25.9 ± 5.3 | [8] |
| 5-HTT* | serotonin | [<sup>11</sup>C]DASB | B<sub>max</sub> | 100 (71) | 25.1 ± 5.8 | [8] |
| 5-HTT* | serotonin | [<sup>11</sup>C]MADAM | BP<sub>ND</sub> | 49 (24) | 39.3 ± 6.4 | [36, 68] |
| CB<sub>1</sub> | cannabinoid | [<sup>18</sup>F]FMPEP-d<sub>2</sub> | V<sub>T</sub> | 20 (0) | 24.4 ± 3.0 | [50] |
| D<sub>2</sub> | dopamine | [<sup>11</sup>C]raclopride | BP<sub>ND</sub> | 16 (7) | 32.7 ± 8.8 | [20] |
| D<sub>2</sub> | dopamine | [<sup>11</sup>C]raclopride | BP<sub>ND</sub> | 47 (0) | 23.5 ± 2.5 | [2, 5, 6] |
| D<sub>2</sub> | dopamine | [<sup>18</sup>F]fallypride | BP<sub>ND</sub> | 49 (33) | 18.4 ± 0.6 | [26] |
| GABA<sub>A/α1</sub> | GABA | [<sup>11</sup>C]Ro15 4513 | V<sub>T</sub> | 27 (n/a) | n/a | [12, 39] |
| GABA<sub>A/α5</sub> | GABA | [<sup>11</sup>C]Ro15 4513 | V<sub>T</sub> | 27 (n/a) | n/a | [12, 39] |
| GABA<sub>A/BZ</sub> | GABA | [<sup>11</sup>C]flumazenil | B<sub>max</sub> | 16 (9) | 26.6 ± 8 | [45] |
| NMDA | glutamate | [<sup>18</sup>F]GE-179 | V<sub>T</sub> | 29 (8) | 40.9 ± 12.7 | [18, 19, 38] |
| mGluR<sub>5</sub> | glutamate | [<sup>11</sup>C]ABP688 | BP<sub>ND</sub> | 27 (12) | 54.6 ± 13.4 | [15] |
| mGluR<sub>5</sub> | glutamate | [<sup>11</sup>C]ABP688 | BP<sub>ND</sub> | 73 (48) | 19.9 ± 3.0 | [62] |
| MOR | opioid | [<sup>11</sup>C]carfentanil | BP<sub>ND</sub> | 86 (42) | 35.6 ± 9.9 | [28, 32, 35, 68] |
| VACHT* | acetylcholine | [<sup>18</sup>F]FEOBV | SUVR | 25 (8) | 36.6 ± 9.7 | [54] |

**Figure 1.**
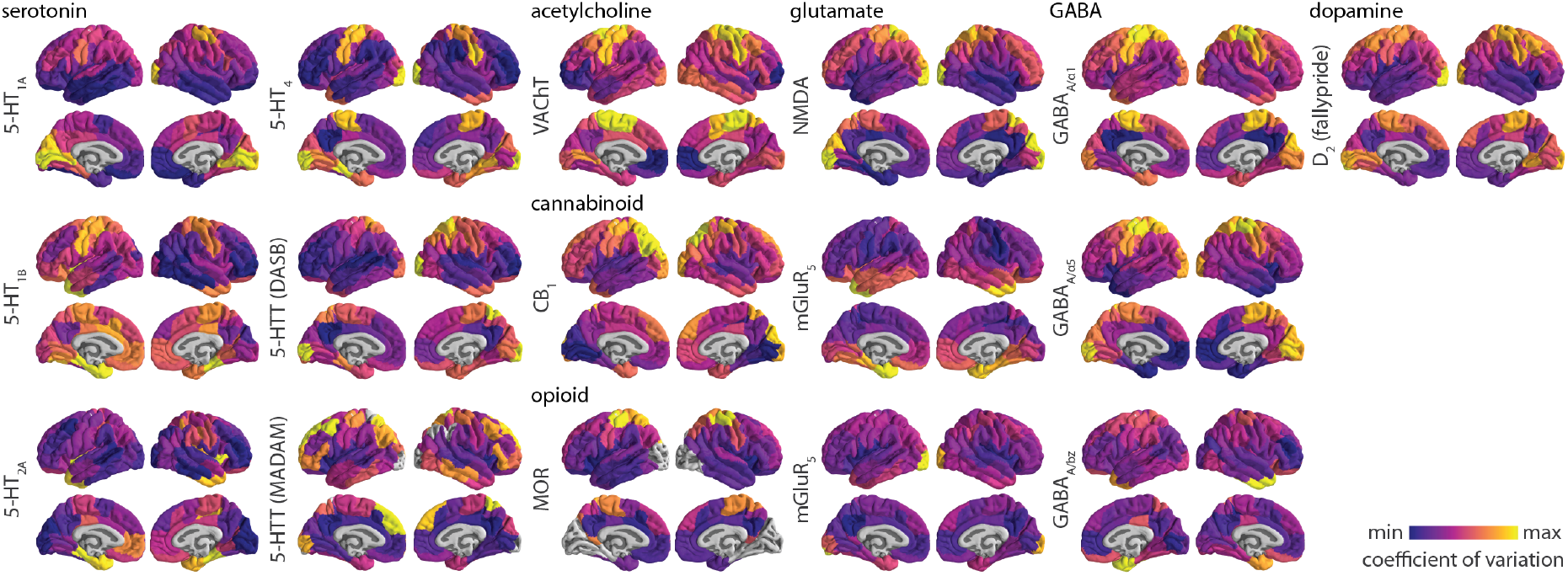
Inter-individual coefficient of variation of receptor/transporter density in the cortex. Inter-individual coefficient of variation is defined as the population standard deviation of tracer binding normalized by population mean, and is calculated for every cortical region. Each coefficient of variation brain map is min-max scaled to showcase the spatial organization of inter-individual variability of neurotransmitter systems. Grey colours reflect regions that have been omitted due to either unstable coefficient of variation or tracer binding quantification reference regions (see *Methods* for details). Two tracers that map 5-HTT were included; tracer names are written in parentheses. GABA_A_ receptors were mapped according to two different subunits (*α*_1_ and *α*_5_) as well as the benzodiazepine binding site (BZ). D_2_ [^11^C]raclopride tracer data is not shown due to high non-displaceable binding in the cortex.

**Figure 2.**
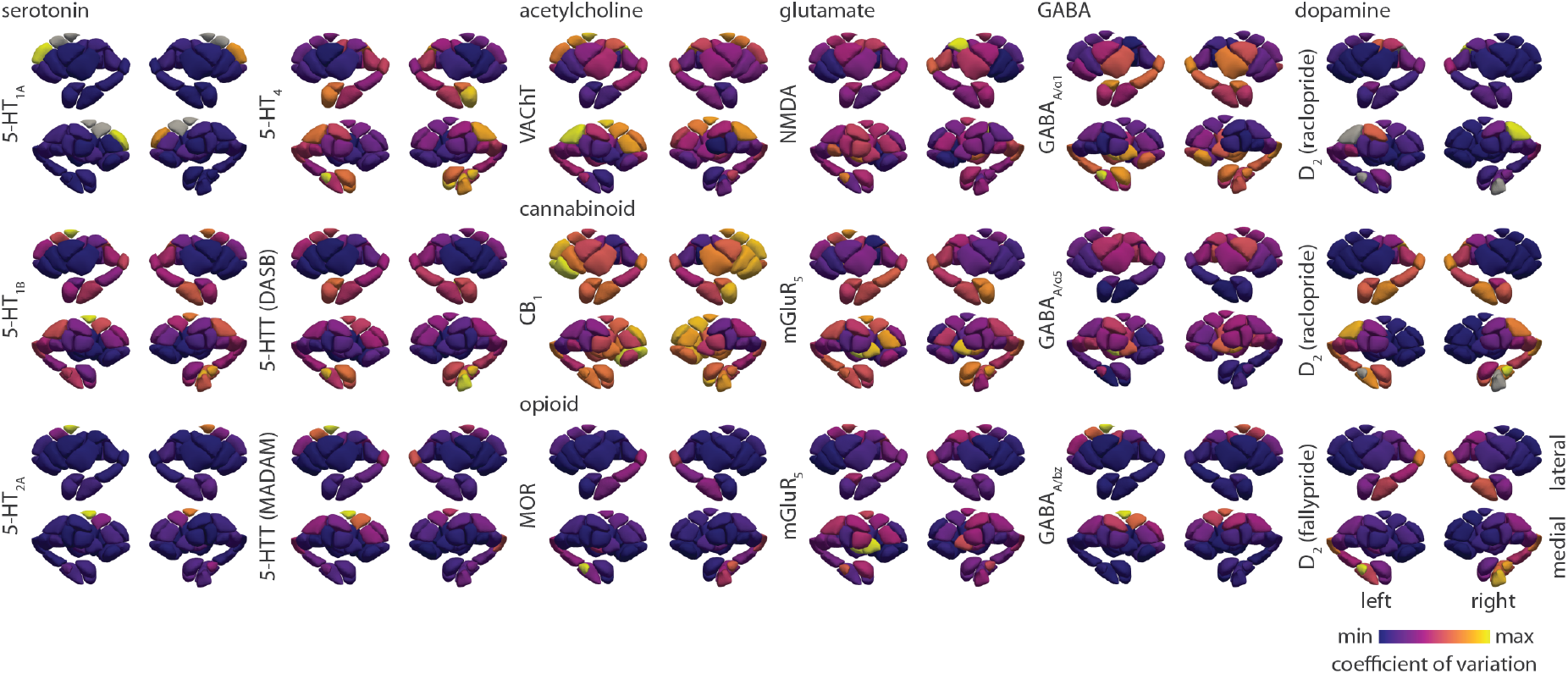
Inter-individual coefficient of variation of receptor/transporter density in the subcortex. Inter-individual coefficient of variation is defined as the population standard deviation of tracer binding normalized by population mean, and is calculated for every subcortical region. Each coefficient of variation brain map is min-max scaled to showcase the spatial organization of inter-individual variability of neurotransmitter systems. Grey colours reflect regions that have been omitted due to unstable coefficient of variation (see *Methods* for details). Tracer names are included in parentheses for 5-HTT and D_2_. GABA_A_ receptors were mapped according to two different subunits (*α*_1_ and *α*_5_) as well as the benzodiazepine binding site (BZ). Note that D_2_ [^11^C]raclopride tracer is only sensitive within the striatum.

In Fig. 3 we show the distribution of cortical and subcortical coefficients of variation for each neurotransmitter receptor and transporter. Density measurements in subcortical structures often vary more than in cortical structures. Within the cortex, inter-individual coefficient of variation is generally low (around 0.2), with some receptors/transporters showing moderate variation (around 0.4, e.g. MOR, CB_1_), and some high variation (*>* 0.5, e.g. NMDA, GABA_A_ *α*_1_ and *α*_5_ subunits). We confirm that the D_2_ tracer [^11^C]raclopride, which is only suitable for quantification of striatal D_2_ receptors [14], shows greatest variation outside of the striatum, as a result of increased measurement noise (Fig. S3).

**Figure 3.**
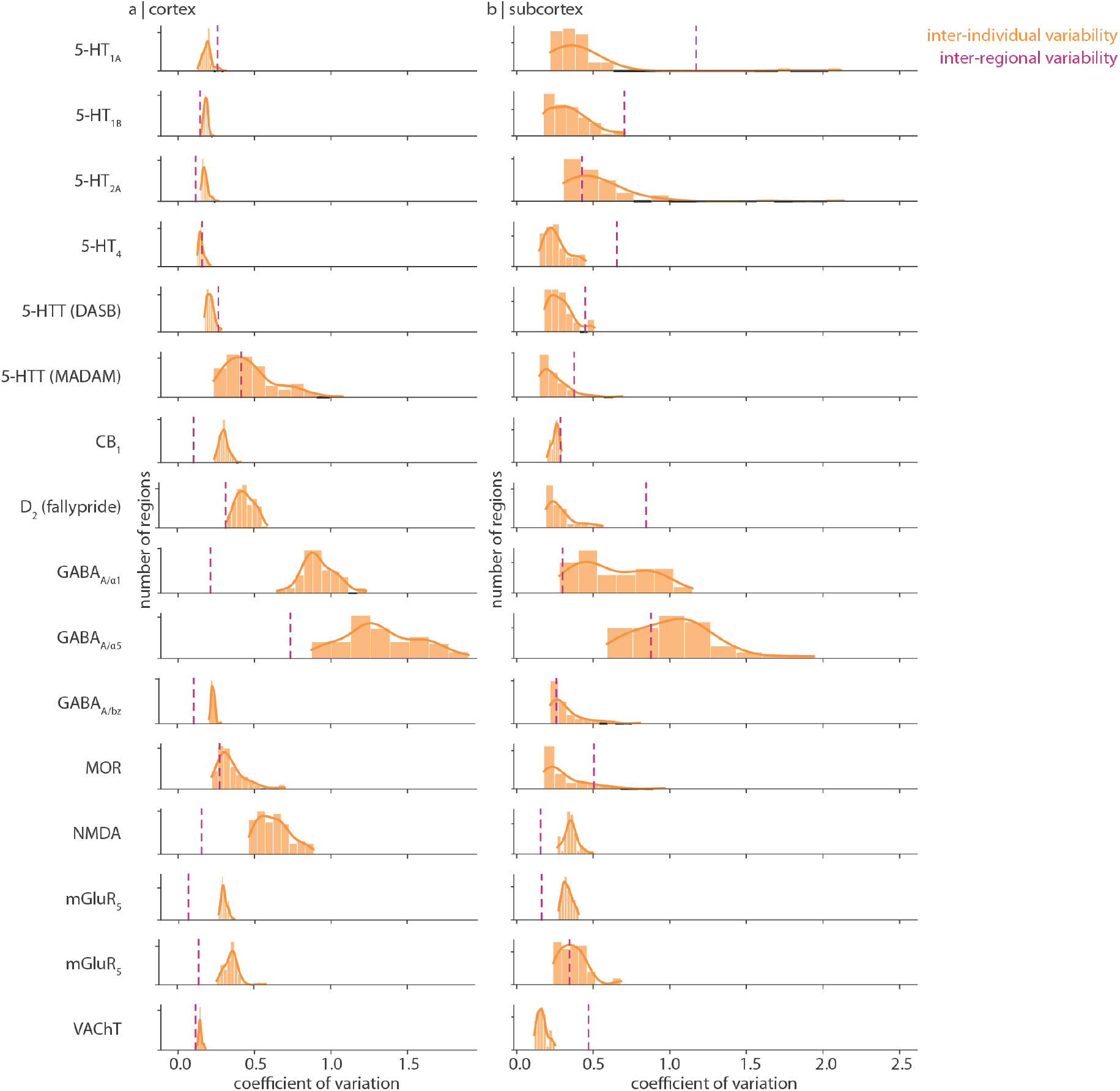
Distributions of inter-individual coefficient of variation. For each receptor and transporter (rows), the distribution of within-region inter-individual coefficient of variation is shown in orange for (a) cortical regions and (b) subcortical regions. These are the same data as shown in Fig. 1 and Fig. 2. A kernel density is estimated for each distribution (solid orange line). The dashed purple line represents the inter-regional coefficient of variation. Coefficient of variation below 0.2 is considered low variability, around 0.5 is moderate variability, and around or above 1 is considered high variability.

In addition, we find that different tracers that bind to the same protein can show different amounts of inter-individual variability, possibly due to differences in study design and preprocessing (e.g. 5-HTT [^11^C]MADAM tracer binding is more variable than 5-HTT [^11^C]DASB tracer binding within the cortex [46, 47]).

Inter-individual variance of a regional measurement is better interpreted in light of the receptor/transporter’s measurement variability across brain regions. To develop this point further, consider a group-averaged measurement with low variation across brain regions (i.e. is approximately homogeneously expressed in the brain) but high variation across individuals. This measurement will have a highly variable spatial profile (i.e. brain map) from one individual to the next. On the other hand, if a measurement varies more across regions than individuals, the regional rank order of protein density will remain similar in all individuals; that is, this measurement will be consistently spatially expressed across individuals. To quantify receptor/transporter density variability across regions, we calculate inter-regional coefficient of variation: the standard deviation of group-averaged receptor/transporter density across brain regions normalized by the mean (Fig. 3 dashed vertical lines; see also schematics in Fig. 4a–c). We find that, in the cortex, many receptors/transporters show similar or greater variability across individuals than regions. Within the subcortex however, receptor/transporter density often varies less across individuals than across regions. This suggests that, although population variance is generally greater in subcortex than in cortex (Fig. 3 yellow bars), subcortical receptor/transporter expression is likely to be stably spatially expressed. Indeed, we find that the ratio of spatial variation to population variation is positively correlated with the out-of-sample consistency of a receptor/transporter’s spatial distribution (i.e. mean pairwise Spearman correlation of receptor/transporter brain maps from different cohorts. *r* = 0.49, *p* = 0.057 within cortex; *r* = 0.77, *p* ≈ 0 within subcortex; Fig. 4). In the cortex, some exceptions to this relationship include glutamatergic mGluR_5_ and endocannabinoid CB_1_, both of which demonstrate highly replicable spatial patterns but low regional-to-population coefficient of variation ratio.

**Figure 4.**
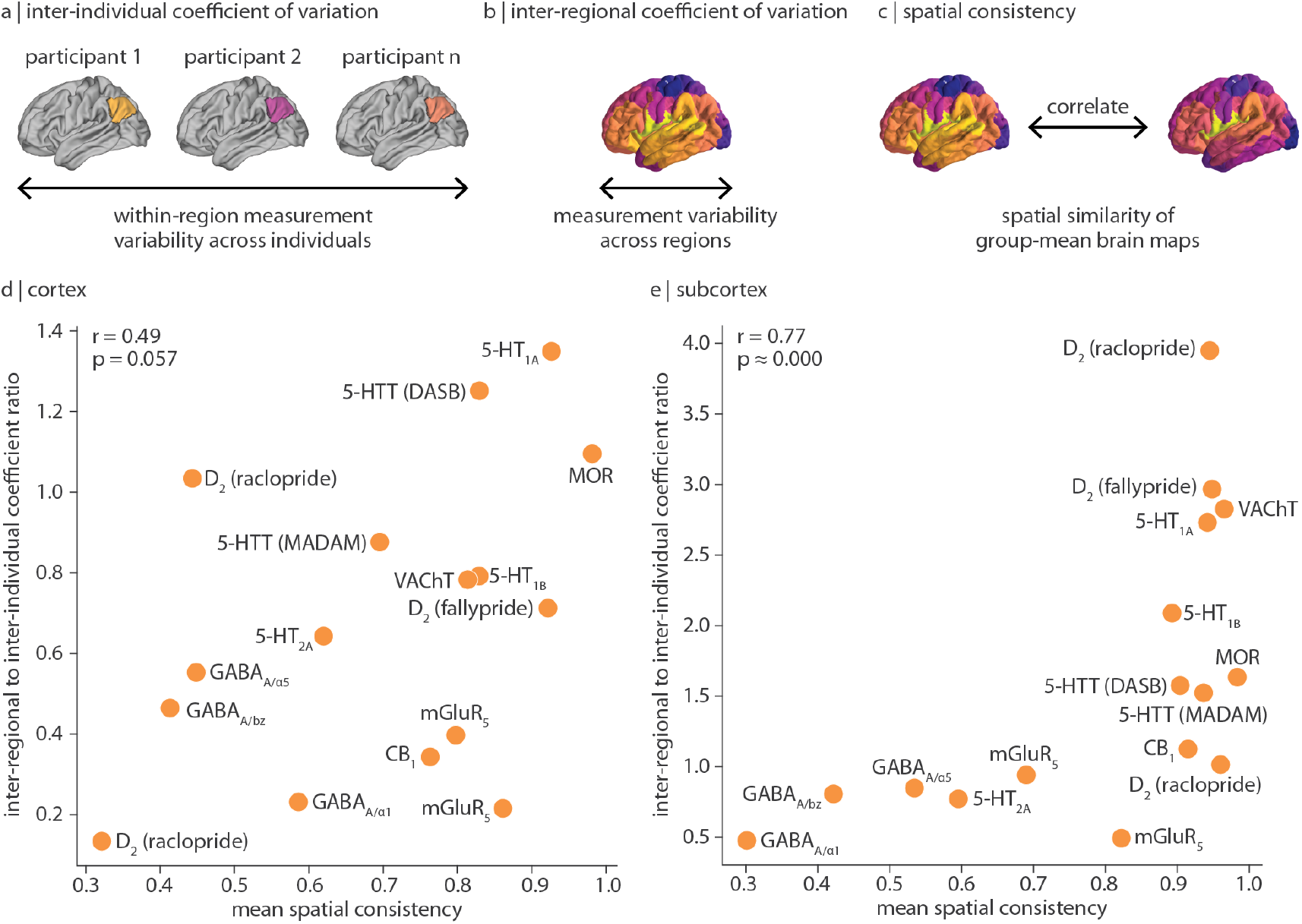
Comparing inter-regional and inter-individual variation of receptor/transporter density. A schematic illustrating three perspectives of variability: (a) inter-individual coefficient of variation quantifies within-region measurement variability across participants; (b) inter-regional coefficient of variation quantifies variability of group-averaged measurements across brain regions; and (c) spatial consistency quantifies the similarity of group-averaged measurements of the same receptor/transporter. For (d) cortex and (e) subcortex, regional-to-population coefficient of variation ratio (*y*-axis) is defined as the inter-regional coefficient of variation (dashed purple line in Fig. 3) normalized by the mean inter-individual coefficient of variation (mean of orange bars in Fig. 3). Values above 1 represent receptors/transporters that vary more across regions than across individuals, and vice versa for values below 1. Note that *y*-axis limits are different in panels (d) and (e). Next, mean spatial consistency is defined as the mean pairwise spatial Spearman’s correlation of group-average tracer images of the same receptor/transporter (*x*-axis). Tracers used for each out-of-sample comparison are detailed in Table S1. Note that GABA_A_ images map different subunits of the GABA_A_ receptor—these receptor subtypes demonstrate unique expression profiles, resulting in lower spatial consistency [61].

## DISCUSSION

In the present report, we estimate standard deviation maps for 12 unique neurotransmitter receptors and transporters to better understand how receptor and transporter density varies across individuals. We show that receptor and transporter variability is heterogeneous across brain regions and systems. Cortical receptor/transporter density typically varies more across individuals than across brain regions, while subcortical receptor/transporter density typically varies less across individuals than across regions. Finally, we show that receptors/transporters that vary more across regions than individuals are also more consistently spatially mapped.

The recent proliferation of group-averaged “reference” brain maps make it possible to spatially relate diverse brain phenotypes with one another [22, 23, 37]. However, the interpretation of such associations is dependent on the generalizability and reliability of these reference maps, which are rarely accompanied by estimates of inter-individual variability [59]. Here we aim to rectify this limitation by retroactively compiling standard deviation maps for previously shared mean receptor density brain maps (see Hansen et al. [23]). We find that inter-individual variability of regional receptor density is organized along specific anatomical landmarks, such that some brain areas vary more across people than others. Surprisingly, while inter-individual variability of structural and functional cortical features is generally greater in transmodal cortex and lower in unimodal cortex [13, 25, 31, 43, 51], we find that the opposite is true for many neurotransmitter receptors and transporters (Fig. 1). As brain maps of inter-individual variability are generated and shared [31, 40, 64], we will better understand how variability varies across brain regions and biological systems.

By combining evidence from multiple lines of analysis, we are able to generate hypotheses regarding the source of variability (e.g. measurement or biological) of different receptors’ expression. In this manuscript, we consider inter-individual variability of regional receptor density measurements as well as out-of-sample spatial consistency in other PET imaging cohorts. However, we can augment our interpretation with reported findings that test out-of-sample spatial replicability using other measurements techniques (e.g. autoradiography, as shown in [8, 21, 45]) and proxies of receptor abundance (e.g. gene expression, as shown in [21, 44, 53]). Take for example serotonergic 5-HT_1A_ density: this receptor is stably expressed across both brain regions and individuals (coefficient of variation around 0.2), spatially replicable across both PET (*r >* 0.9) and autoradiography (*r >* 0.6) cohorts, and strongly correlated with its protein-coding gene (*r* = 0.88), indicating a protein with approximately the same regional receptor abundance in any brain (i.e. low biological variability, low measurement variability, and conserved spatial expression) [8, 21, 23, 44]. Similarly, the endocannabinoid receptor CB_1_ and opioid receptor MOR demonstrate spatial consistency (mean *r >* 0.75) and high coexpression with their protein-coding genes (*CNR1* (*r* = 0.74) and *OPRM1* (*r* = 0.84) respectively, as reported in [21]). However, their regional receptor abundance is variable across people (coefficient of variation around 0.4). This suggests that, while the spatial distributions of these proteins are consistent, they may exhibit an individual-specific baseline shift (i.e. high biological variability, low measurement variability, and conserved spatial expression). Finally, there are receptors that are systematically inconsistently expressed. Ionotropic (and heteromeric) receptors GABA_A_ (*α*_1_ and *α*_5_ subunits) and NMDA show high population variability in regional receptor abundance (coefficient of variation *>* 0.5) and GABA_A_’s spatial patterning is only moderately replicable in separate PET (*r* ≈ 0.5) and autoradiography (*r* = 0.20) cohorts. Such inconsistent measurements may reflect noise [58], individual-specific expression [4, 29, 42], protein turnover rate (i.e. temporal variability), or individual differences in receptor subunit composition.

We end with a note on interpretation. First, while we show brain maps of inter-individual coefficient of variation in the cortex and subcortex (Fig. 1, 2), these maps are min-max scaled and in many cases (e.g. the serotonergic receptors), the inter-individual coefficient of variation is consistently very low. Fig. 3 should be used to compare the variability across tracers. Second, our measurement of inter-individual variability is agnostic to whether the source of variability is individual differences, measurement noise, or study design (e.g. modelling technique) [47]. To better assess the generalizability and replicability of receptor brain maps, we apply our own out-of-sample comparisons and we draw on our earlier work comparing alternative PET tracers, imaging modalities, and protein-coding gene expression [21, 23]. Third, due to ethical restrictions in sharing individual data, we are unable to test whether receptor binding is normally distributed across individuals. Individual outliers may therefore skew the standard deviation.

In summary, we assemble an atlas of neurotransmitter receptor and transporter density variability. This atlas complements our previously published atlas of wholebrain receptor/transporter densities [23]. Our work sheds light on how receptor systems vary in healthy individuals, and provides a means of assessing the generalizability of PET-derived receptor density quantification.

## METHODS

All code and data used to conduct the analyses are available at https://github.com/netneurolab/hansen_receptorvar.

### PET data acquisition

Our group had previously assembled group-averaged PET tracer images for 19 neurotransmitter receptors and transporters from research groups and PET imaging centers globally [23]. In an effort to better understand how these measurements vary across individuals, we recontacted all collaborators who had contributed mean receptor maps and asked whether they would be interested in providing group mean and standard deviation images for each tracer. Altogether we compiled 18 tracer mean and standard deviation images, encompassing 12 unique neurotransmitter receptors and transporters, and 7 neurotransmitter systems. Each study, the associated receptor/transporter, tracer, number of healthy participants, age, and reference with full methodological details of data acquisition can be found in Table 1. In all cases, only scans from healthy participants were included. Group mean and standard deviation images were registered to MNI152NLin6Asym space, then parcellated according to 100 cortical regions as defined by the Schaefer parcellation [57] and 54 subcortical regions as defined by the Melbourne Subcortex Atlas S4 [66].

We note some tracer-specific special cases: (1) while tracer binding for most neurotransmitter receptors is estimated using the cerebellum as the reference region, the mu-opioid receptor (MOR) is measured using the occipital cortex as the reference region. We therefore set all regions in the occipital cortex to NaN. (2) Three dopaminergic D_2_ images were shared, two measured with the tracer [^11^C]raclopride and one measured with the tracer [^18^F]fallypride. Due to the lower affinity of [^11^C]raclopride to D_2_ receptors, this tracer can only reliably estimate binding in regions with high D_2_ density (i.e. the striatum) [49]. [^11^C]raclopride measurements outside of the striatum are therefore expected to demonstrate large variation across participants. On the other hand, [^18^F]fallypride is primarily suitable for estimation of extra-striatal D_2_ receptors [26, 70]. (3) Two serotonergic 5-HTT images acquired using different tracers ([^11^C]DASB and [^11^C]MADAM) were shared. We include both for comparison. (4) Two subunits (*α*_1_ and *α*_5_) of the GABA_A_ receptor were mapped using a single PET tracer [^11^C]Ro15-4513 by way of spectral analysis [39]; we include both for comparison. We also include [^11^C]flumazenil, a tracer that binds to the benzodiazepine (BZ) binding site of GABA_A_ receptors [45]. Although subunits *α*_1_, *α*_5_, and benzodiazapine are all part of the GABA_A_ receptor, they demonstrate diverse spatial profiles [61]. (5) Two mGluR_5_ images were shared, both measured using [^11^C]ABP688; we include both for comparison.

Finally, to estimate the spatial consistency of receptor/transporter density maps, we calculate the average spatial correlation between a receptor’s mean tracer image with any other mean tracer image for this receptor, both from within the set of maps analyzed here, and from out-of-sample mean tracer images from the previously mentioned PET receptor atlas [23]. In other words, for each receptor/transporter, we calculate 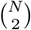 correlations (where *N* is the number of mean tracer images available for this receptor/transporter), then calculate their average. See Table S1 for a complete list of images that were correlated with each receptor/transporter density map. Note that out-of-sample mean receptor density maps are not accompanied by standard deviation maps, and they may be collected using a different PET tracer. Furthermore, all MOR [^11^C]carfentanil images were collected at the same PET centre and group maps may not be independent. Mean spatial consistency for MOR is therefore likely inflated.

### Coefficient of variation

In biological systems, the standard deviation of a distribution of measurements typically scales with the mean (see also Fig. S1 and Fig. S2). Therefore, rather than directly analyzing standard deviation values, we normalized the standard deviation by the mean. This ratio is called the coefficient of variation. In this work, we consider the coefficient of variation of tracer binding measurements (i.e. neurotransmitter receptor/transporter densities) both across individuals (“inter-indivudal”) and across regions (“inter-regional”). When calculated across individuals, there is one coefficient of variation value per region, representing inter-individual variability of within-region receptor/transporter density. The coefficient of variation can be unstable when the mean (denominator) approaches 0. Therefore, when calculating coefficient of variation, we omit the regions whose mean tracer binding is in the bottom fifth percentile, if tracer binding values are below 0.1.

Likewise, when calculated across regions rather than individuals, there is one inter-regional coefficient of variation value per brain map, representing how much receptor/transporter density varies across brain regions. More specifically, the standard deviation of mean tracer binding across brain regions (for cortex and subcortex separately) is divided by the mean tracer binding across brain regions. Finally, the regional-to-population coefficient of variation ratio is calculated as the inter-regional coefficient of variation divided by the mean inter-individual coefficient of variation. Values above 1 reflect neurotransmitter receptors/transporters that vary more across brain regions than across individuals, and values below 1 reflect neurotransmitter receptors/transporters that vary more across individuals than brain regions.

## Acknowledgments

BM acknowledges support from the Natural Sciences and Engineering Research Council of Canada (NSERC), Canadian Institutes of Health Research (CIHR), Brain Canada Foundation Future Leaders Fund, the Canada Research Chairs Program, the Michael J. Fox Foundation, and the Healthy Brains for Healthy Lives initiative. JYH acknowledges support from the Helmholtz International BigBrain Analytics & Learning Laboratory, NSERC, and CIHR. SG and LT acknowledge support from CIHR.

---

**Figure S1.**
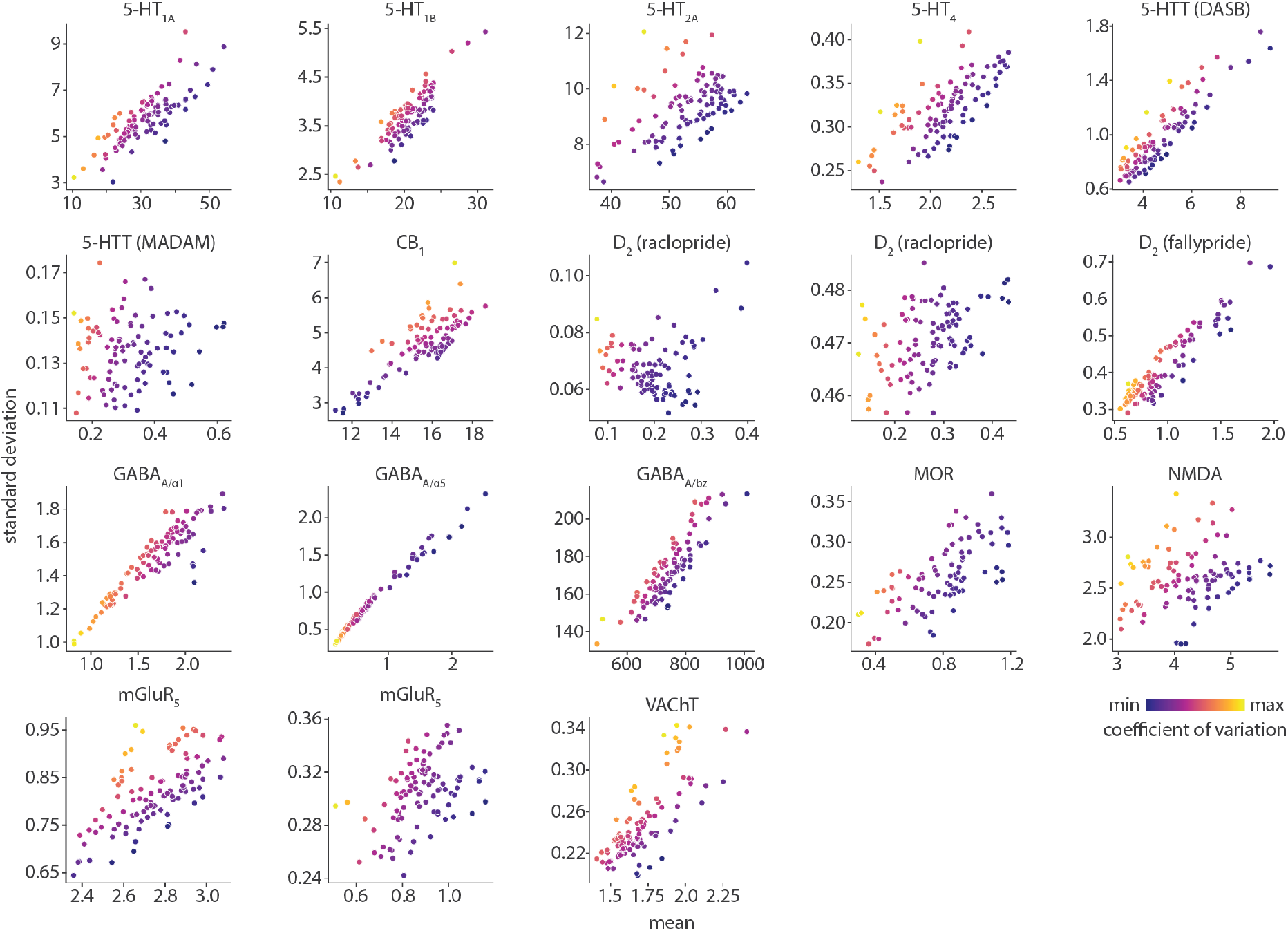
Correlation between mean and standard deviation of receptor/transporter density within cortex. Mean tracer binding (*x*-axis) is correlated with standard deviation of tracer binding (*y*-axis) across individuals. Each circle is a cortical region (*n* = 100). Circle colour represents inter-individual coefficient of variation (as shown in Fig. 1).

**Figure S2.**
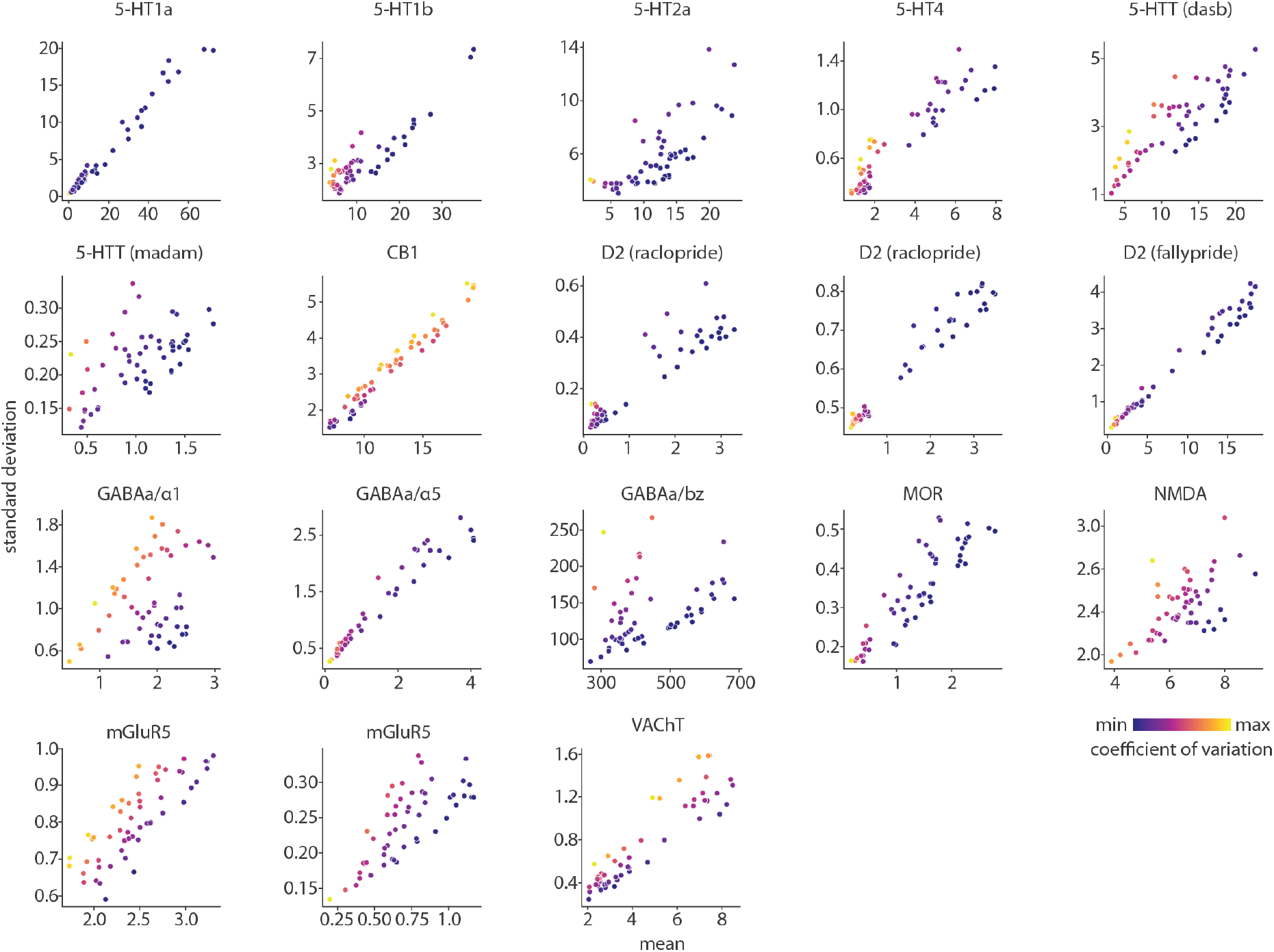
Correlation between mean and standard deviation of receptor/transporter density within subcortex. Mean tracer binding (*x*-axis) is correlated with standard deviation of tracer binding (*y*-axis) across individuals. Each circle is a subcortical region (*n* = 54). Circle colour represents inter-individual coefficient of variation (as shown in Fig. 2).

**Figure S3.**
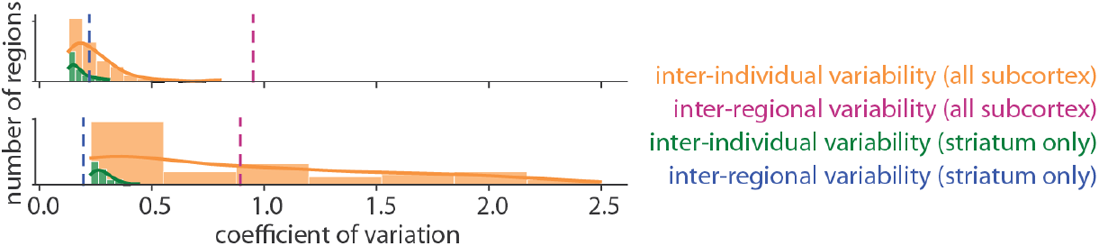
Subcortical distributions of inter-individual coefficient of variation for D_2_ [^11^C]raclopride tracer. We show the distribution of subcortical inter-individual coefficient of variation (orange) and striatal inter-individual coefficient of variation (green). A kernel density is estimated for each distribution (solid lines). The dashed purple line represents the inter-regional coefficient of variation across all subcortical structures, and the dashed blue line represents inter-regional coefficient of variation across all striatal regions. Notably, variability is considerably lower in the striatum where [^11^C]raclopride tracer is sensitive to D_2_ receptor abundance. Data from [20] (*N* = 16) is shown on the top and data from [2, 5, 6] (*N* = 47) is shown on the bottom.

**TABLE S1.** Out-of-sample group-average receptor/transporter density maps. To calculate mean spatial consistency in Fig. 4, we correlate each receptor and transporter’s mean tracer image (“original map”) with any other available mean tracer image for this receptor/transporter (“other map(s)”), both from within the set of maps analyzed here, and from out-of-sample mean tracer images from the PET receptor atlas introduced in Hansen et al. [23]. Note that MOR [^11^C]carfentanil maps were pulled from the same centre and therefore group maps are not necessarily independent.

| Receptor/Transporter | Original map (*N*) | Other map(s) (*N*) |
| --- | --- | --- |
| 5-HT<sub>1A</sub> | [<sup>11</sup>C]CUMI-101 (8) [8] | [<sup>11</sup>C]WAY-100635 (35) [56] |
| 5-HT<sub>1B</sub> | [<sup>11</sup>C]AZ10419369 (36) [8] | [<sup>11</sup>C]P943 (23) [56]<br>[<sup>11</sup>C]P943 (65) [17] |
| 5-HT<sub>2A</sub> | [<sup>11</sup>C]Cimbi-36 (29) [8] | [<sup>18</sup>F]Altanserin (19) [56]<br>[<sup>18</sup>F]MDL100907 (3) [65] |
| 5-HTT | [<sup>11</sup>C]DASB (100) [8] | [<sup>11</sup>C]DASB (30) [56]<br>[<sup>11</sup>C]MADAM (49) [36, 68] |
| 5-HTT | [<sup>11</sup>C]MADAM (49) [36, 68] | [<sup>11</sup>C]DASB (30) [56]<br>[<sup>11</sup>C]DASB (100) [8] |
| CB<sub>1</sub> | [<sup>18</sup>F]FMPEP-d<sub>2</sub> (20) [50] | [<sup>18</sup>F]FMPEP-d<sub>2</sub> (22) [33]<br>[<sup>11</sup>C]OMAR (77) [48] |
| D<sub>2</sub> | [<sup>11</sup>C]raclopride (16) [20] | [<sup>11</sup>C]raclopride (47) [2, 5, 6]<br>[<sup>11</sup>C]raclopride (7) [3]<br>[<sup>18</sup>F]fallypride (49) [26]<br>[<sup>18</sup>F]FLB457 (37) [63]<br>[<sup>18</sup>F]FLB457 (55) [55] |
| D<sub>2</sub> | [<sup>11</sup>C]raclopride (47) [2, 5, 6] | [<sup>11</sup>C]raclopride (16) [20]<br>[<sup>11</sup>C]raclopride (7) [3]<br>[<sup>18</sup>F]fallypride (49) [26]<br>[<sup>18</sup>F]FLB457 (37) [63]<br>[<sup>18</sup>F]FLB457 (55) [55] |
| D<sub>2</sub> | [<sup>18</sup>F]fallypride (49) [26] | [<sup>18</sup>F]FLB457 (37) [63]<br>[<sup>18</sup>F]FLB457 (55) [55] |
| GABA<sub>A</sub>/α<sub>1</sub> | [<sup>11</sup>C]Ro154513 (27; α<sub>1</sub>) [39] | [<sup>11</sup>C]Ro154513 (27; α<sub>5</sub>) [39]<br>[<sup>11</sup>C]flumazenil (16; BZ) [45] |
| GABA<sub>A</sub>/α<sub>5</sub> | [<sup>11</sup>C]Ro154513 (27; α<sub>5</sub>) [39] | [<sup>11</sup>C]Ro154513 (27; α<sub>1</sub>) [39]<br>[<sup>11</sup>C]flumazenil (16; BZ) [45] |
| GABA<sub>A</sub>/BZ | [<sup>11</sup>C]flumazenil (16; BZ) [45] | [<sup>11</sup>C]Ro154513 (27; α<sub>1</sub>) [39]<br>[<sup>11</sup>C]Ro154513 (27; α<sub>5</sub>) [39] |
| mGluR<sub>5</sub> | [<sup>11</sup>C]ABP688 (27) [15] | [<sup>11</sup>C]ABP688 (73) [62]<br>[<sup>11</sup>C]ABP688 (22) [23] |
| mGluR<sub>5</sub> | [<sup>11</sup>C]ABP688 (73) [62] | [<sup>11</sup>C]ABP688 (27) [15]<br>[<sup>11</sup>C]ABP688 (22) [23] |
| MOR | [<sup>11</sup>C]carfentanil (86) [28, 32, 35, 68] | [<sup>11</sup>C]carfentanil (204) [30]<br>[<sup>11</sup>C]carfentanil (39) [69] |
| VACHT | [<sup>18</sup>F]FEOBV (25) [54] | [<sup>18</sup>F]FEOBV (5) [7]<br>[<sup>18</sup>F]FEOBV (18) [1] |

